# Multi-omics analysis reveals genetic regulatory mechanisms of muscle traits in wild and domesticated Bactrian camels

**DOI:** 10.1101/2025.09.02.673668

**Authors:** Baojun De, Bowen Lei, Yuting Chen, Lu Li, Yiyi Liu, Fanhua Meng, Ting Jia, Shenyuan Wang, Tao Li, Chunxia Liu, Hongmei Xiao, Fang Wan, Yingchun Liu, Huanmin Zhou, Wenguang Zhang, Bing Han, Xin Wen, Hai Long, A Naer, Hongmei Bao, Qian Song, Ha Sichaolu, Yanru Zhang, Yuwen Liu, Junwei Cao

## Abstract

Bactrian camels, large domesticated animals adapted to extreme environments, have evolved under natural and artificial selection to suit their living conditions and meet human needs. In arid and semi-arid regions, they provide valuable resources through economically important traits such as meat, milk, and wool production, contributing significantly to both economy and ecology. However, research on gene function, adaptive evolution, and molecular mechanisms underlying complex traits in Bactrian camels remains limited, constraining the full utilization of their breeding potential and conservation efforts.To address this gap, we performed whole-genome resequencing (WGS) on 35 domesticated Bactrian camels from the Chinese Sunite local population and integrated this data with publicly available WGS data from 112 Asian Bactrian camels. Our comprehensive analysis revealed significant genetic differences and genomic features between wild and domesticated Bactrian camels. We further integrated 27 epigenomic (ATAC-seq) and transcriptomic (RNA-seq) datasets from eight biologically important tissues of four Bactrian camels, along with skeletal muscle scRNA-seq and scATAC data. This allowed us to identify tissue-specific genes, regulatory elements, and genes under positive selection.Our integrated analysis identified target genes and their upstream regulatory elements, including potential regulatory transcription factors, with a focus on positively selected regions in Bactrian camel muscle tissue. We constructed regulatory pathways from key SNPs to regulatory elements to genes, and for the first time, identified regulatory elements in major tissues of the Bactrian camel genome. Our results also summarized the single-cell transcriptional landscape of Bactrian camel skeletal muscle, revealing cellular heterogeneity and genetic expression patterns across two developmental stages.This comprehensive regulatory element map provides a valuable resource for genetic and genomic studies in Bactrian camels. It reveals key regulatory networks, important genes, and regulatory elements influencing muscle development, as well as potentially significant functional variants affecting meat production traits. These findings will provide crucial data support for future genetic improvement and breeding efforts in Bactrian camels.

## 2 Introduction

Camels play an irreplaceable role in food security in arid and semi-arid regions of Africa and Asia. Camel meat is valued for its low fat and cholesterol content, providing not only a nutritious option but also showing potential in treating various ailments including seasonal fever, sciatica, shoulder pain and asthma^[1]^. The high nutritional value, low allergenicity and presence of small nanobodies in camel milk demonstrate significant potential in treating multiple diseases, including diabetes and cancer, garnering widespread attention in health and immunology fields^[2,3,4,5]^. As a research model, camels have shown unique value and broad potential in immune systems and single-domain antibodies^[6]^, Middle East Respiratory Syndrome (MERS) coronavirus^[7]^, and desert adaptation mechanisms^[8]^.

During domestication, artificial selection and climate change-driven natural selection have jointly become key drivers shaping genetic characteristics and phenotypic traits of animal populations globally^[9]^. To date, comparative genomics and whole-genome resequencing have identified genomic variations involved in domestication and genetic improvement of several domesticated animals such as pigs^[10]^, cattle^[11]^, chickens^[12]^, and sheep^[13]^, determining regions and genes under selection in domesticated species. However, functional DNA variations affecting phenotypic variation exist not only within traditional transcriptional regulatory regions but are also widely distributed in non-coding regions beyond regulatory elements^[14]^. Selective sweeps have revealed numerous non-coding regulatory DNA variations outside coding regions, which play crucial roles in driving phenotypic diversity. While coding region variations may have broad effects across multiple tissues, variations in cis-regulatory elements can influence gene expression in specific spatiotemporal patterns, thereby promoting morphological evolution and phenotypic differentiation^[15]^.

Nevertheless, for large livestock, especially Bactrian camels, functional annotation of the non-coding genome remains insufficient, limiting the ability to identify non-coding DNA variations leading to phenotypic differences^[16]^.

Single-cell RNA sequencing (scRNA-seq)^[17]^ technology, compared to traditional RNA-seq analysis, enables precise acquisition and analysis of transcriptome information from individual cells in animal tissues, overcoming the limitation of obtaining only homogenized RNA expression information at the multi-cellular tissue level. By revealing gene expression differences among distinct cells within the same tissue, it has facilitated rapid identification and genetic improvement of desirable traits in livestock and poultry, and deepened our understanding of cell-specific expression and regulatory mechanisms in biological processes such as growth and development, immune response, and disease resistance^[18,19]^.

Although whole-genome resequencing technology has shown potential in revealing population structure and identifying phenotype-related functional genes involved in domestication and breeding processes of Bactrian camels^[20]^, large-scale multi-tissue transposase-accessible chromatin sequencing (ATAC-seq)^[21]^ and single-cell RNA-seq technologies have not yet been utilized to map and describe open chromatin regions (OCRs) in camels, or to assist in identifying functional genes associated with their key economic traits. Therefore, the integration of multi-omics technologies has become an important approach for comprehensive insights into the biological characteristics of organisms. Through integrated analysis and in-depth data mining, it provides new strategies for genetic breeding of livestock and poultry, contributing to the cultivation of breeds with superior traits.

In this study, we integrated resequencing data from 35 Chinese Sunite local breed Bactrian camels and publicly available genomic data from 112 camels^[22]^ to identify selected gene regions and genome-wide variation sites in Bactrian camels. Using RNA-seq and ATAC-seq technologies, we constructed epigenetic maps for eight major tissues of Bactrian camels, annotated regulatory element functions, and identified transcription factors. Furthermore, we discovered regulatory elements and genes associated with muscle tissue characteristics and analysed regulatory mechanisms of key genes within selected regions. Subsequently, by integrating single-cell RNA sequencing (scRNA-seq) and single-cell ATAC data from adult and juvenile Bactrian camel longissimus dorsi muscles, we explored in depth how domestication and artificial selection influence changes in skeletal muscle cell atlases affecting skeletal muscle phenotype development.These research findings are expected to optimize breeding strategies and provide scientific support for sustainable development in animal husbandry. Concurrently, population resequencing data, multi-tissue chromatin accessibility analysis data, single-cell transcriptome data, and single-cell ATAC data will become valuable resources for Bactrian camel domestication and breeding research.

## 3 METHODS

### 3.1 Sample collection

We selected 35 camels from three farms in Sonid Left Banner and Sonid Right Banner, Xilingol League, Inner Mongolia, China, for whole-genome sequencing studies. From each camel, 50 ml of blood was collected via jugular venipuncture, treated with disinfectant, stored in EDTA anticoagulant tubes, and preserved at −80°C. Skeletal muscle samples from Sunite Bactrian camels were obtained from 6-year-old adult females during lactation and 4-day-old female camels in Sonid Left Banner.

Tissue samples from two developmental stages were collected from four Sunite Bactrian camels, including two newborn female calves and two adult females. Eleven types of tissues were sampled: skeletal muscle, fat, heart, liver, spleen, cerebrum, cerebellum, kidney, rumen, reticulum, and omasum. These tissue samples underwent sequencing analysis using techniques such as RNA-seq and ATAC-seq to identify novel transcripts and cis-regulatory sequences in the Sunite Bactrian camel genome, including open chromatin regions, enhancers, and promoters.

All Bactrian camels were fasted the night before humane euthanasia. The care and slaughter of experimental animals complied with animal welfare laws, guidelines, and policies.

### 3.2 Whole-genome resequencing

#### Mapping and assembly

For processing the resequencing data of 35 camels, we first performed quality control using Fastp 0.23.1^[23]^, removing reads with quality scores below 20 and sequences shorter than 35 bases to ensure high-quality data of appropriate size. Subsequently, we aligned the preprocessed data to the camel genome (using the NCBI GCF_009834535.1 BCGSAC_Cfer_1.0 assembly) using the “bwa mem” command in BWA 0.7.17. The sequences were then sorted using SAMTools 1.15.1^[24]^ to ensure data integrity.

Next, we used the “MarkDuplicates” command in Picard 3.0^[25]^to mark and remove duplicates introduced by PCR. We then employed the “HaplotypeCaller” command in GATK 4.2.3.0-0^[26]^ to detect variant information on different chromosomes for each sample, generating GVCF files. The GVCF files for all individuals on the same chromosome were merged using the “CombineGVCFs” command, followed by genotype analysis using the “GenotypeGVCFs” command. Finally, we integrated the results using the “MergeVcfs” command to produce a raw VCF file containing genetic variation information for each sample, which was used for subsequent bioinformatic analyses.This entire workflow aims to extract genetic variation information from the original sequencing data of camel individuals, providing a foundation for further bioinformatic research.

#### Filtering and quality control

We applied a series of filtering steps using the VariantFiltration function in GATK to eliminate low-quality SNP sites. Specifically, we employed the following filtering criteria: QD < 2.0, MQ < 40.0, FS > 60.0, SOR > 3.0, and MQRankSum < −12.5. Subsequently, we performed additional filtering steps, including the removal of sites with MAF < 0.05, elimination of multi-allelic sites, exclusion of sex chromosome loci, removal of highly correlated samples, and exclusion of SNP samples from undetermined breeds to ensure data quality and consistency.

### 3.3 Variant calling

Raw sequencing data underwent quality control using Fastp v0.23.2, removing adapter sequences, bases with quality below 15, reads shorter than 15 bp, and reads containing more than 5 unknown bases (N). Subsequently, quality-controlled reads from each sample were aligned to the Bactrian camel reference genome (https://ftp.ncbi.nlm.nih.gov/genomes/all/GCF/009/834/535/GCF_009834535.1_BCGSAC_Cfer_1.0/GCF_009834535.1_BCGSAC_Cfer_1.0_genomic.fna.gz) using BWA-MEM v0.7.17. SAMtools v1.6²⁷ was then used to convert SAM format to BAM format. PCR duplicates were removed using Picard v2.27.5, and GATK v4.2.6.1 was employed for base quality recalibration and variant calling.

Raw variants were filtered based on the following criteria: minor allele frequency < 5%, call rate < 90%, removal of sex chromosomes and mitochondrial chromosomes, and elimination of multi-allelic sites. After filtering, 6,208,574 SNPs and 928,685 Indels were retained for subsequent analyses.

### 3.4 Population genetic structure

Population genetic structure was analysed using 6,208,574 SNPs from 146 camels. Principal component analysis (PCA) was performed using the “--pca” function in Plink v1.9 based on the variance-standardized relationship matrix for 14 populations. The proportion of variance explained by each principal component was calculated as the ratio of its corresponding eigenvalue to the sum of all eigenvalues.A phylogenetic tree was constructed using the neighbour-joining method based on a distance matrix calculated by VCF2Dis v1.47. ADMIXTURE v1.3.0 was used to infer ancestral components of populations using maximum likelihood estimation. The optimal number of ancestral populations (K) was evaluated by calculating cross-validation error rates for K values ranging from 1 to 14.Migration events between camel populations were inferred using TreeMix v1.13^[28]^. The analysis was performed with migration edges (m) ranging from 1 to 10, using the parameter “-k 500”. Maximum likelihood trees were constructed, and corresponding residual matrices were generated.

### 3.5 Genetic diversity and differentiation

Genomic heterozygosity for each population was calculated using the “--hardy” function in Plink v1.9. Nucleotide diversity (π) and population differentiation (F-statistics, Fst) were computed for each population using VCFtools v0.1.16^[29]^ in sliding windows of 50 kb with a step size of 25 kb. Genome-wide linkage disequilibrium patterns for each population were assessed using PopLDdecay v3.42, with the maximum distance for linkage set to 500 kb.

### 3.6 Evolutionary selection among populations

Selection signatures between Sunite and wild camels were detected using three methods: population differentiation (F-statistics, Fst), nucleotide diversity (π), and cross-population extended haplotype homozygosity (XP-EHH). VCFtools v0.1.16 and the rehh package were used for these analyses. The genome was divided into sliding windows of 50 kb with a step size of 25 kb. Regions in the top 1% of each statistic were considered as candidates under strong positive selection. Overlapping regions identified by at least two methods were subjected to Kyoto Encyclopedia of Genes and Genomes (KEGG)^[30]^and Gene Ontology (GO)^[31]^ enrichment analyses. Enriched terms with P-value < 0.05 were considered significant.

### 3.7 RNA-seq library preparation and sequencing

Total RNA was extracted using TRIzol reagent (Thermo Scientific, 15596026) from skeletal muscle, mammary gland, rumen, reticulum, and omasum of adult and juvenile Bactrian camels, and corresponding tissues (plus liver) of dromedary camels. RNA quality was assessed by RIN values (range 6.10–9.00, 90% samples >7.00). After rRNA depletion, strand-specific RNA-seq libraries were constructed following Illumina protocols.Sequencing data were preprocessed using trim_galore (v0.6.6) for adapter trimming and low-quality read removal (Q<20). Clean reads were aligned to the reference genome BCGSAC_Cfer_1.0 using hisat2 (v2.2.1) with “--rna-strandness RF” parameter. Aligned BAM files were counted using featureCounts (v2.0.1, s=2). Differential expression analysis was performed using DESeq2^[32]^, with spike-in normalization factors and an experimental design accounting for developmental stage.Tissue-specific genes were identified based on an expression threshold of TPM>1, normalized using the qsmooth package, and Z-scores calculated. Genes with expression at least three-fold higher than in other tissue samples were defined as tissue-specific. Gene co-expression network analysis was conducted using the WGCNA package to evaluate co-expressed gene sets associated with camel tissues and developmental ages.Gene Ontology enrichment analysis for biological processes was performed using clusterProfiler and org.Hs.eg.db packages. Hypergeometric distribution tests were used to assess the significance of differentially expressed gene enrichment in selected gene sets from co-expression analysis.

### 3.8 ATAC-seq library preparation and sequencing

Approximately 5 mg of tissue from skeletal muscle, mammary gland, liver, rumen, reticulum, and omasum samples were pulverized in liquid nitrogen. The pulverized tissue was suspended in 1 mL of ice-cold PBS, pelleted, and resuspended in 1 mL of lysis buffer (50 mM HEPES pH 7.5, 140 mM NaCl, 1 mM EDTA pH 8.0, 10% glycerol, 0.5% NP-40, 0.25% Triton X-100).

Data processing followed these steps: adapter trimming using trim_galore (v0.6.6), alignment to the camel reference genome using Bowtie2 (v2.4.5), filtering with SAMTools (v1.15.1) and Sambamba (v0.8.2), and peak calling using MACS3 (v3.0.0a7). Quality control included assessment of NSC, RSC, and FRiP scores, and calculation of TSS enrichment scores using ATACseqQC (v1.20.2).

For cis-regulatory element analysis, promoters and distal enhancers were defined. Motif enrichment analysis on tissue-specific elements was performed using HOMER’s findMotifsGenome.pl (v4.1). Element enrichment in specific regions was analyzed using the bedtools shuffle function from BEDTools, with p-values and fold changes (FC) calculated.

### 3.9 scRNA-seq data processing

All samples were submitted and sequenced on the same day to minimize batch effects. Sample preparation was performed using the 5’ assay on the 10X Chromium platform (10X Genomics), targeting 20,000 cells per diet group (approximately 5,000 cells per biological replicate). Overall, PE-150 sequencing was conducted on an Illumina NovaSeq6000, aiming for 50,000 reads per cell. Sample processing and sequencing were completed in a single run at the VANderbilt Technologies for Advanced GEnomics core (VANTAGE).10× Genomics scRNA-seq raw data were processed and aligned to the reference genome downloaded from ENSEMBL (release 104), then quantified using the Cell Ranger software package (version 6.0.2). Downstream analysis was performed using the R package Seurat (version 4.1.3)^[33]^. Cells with low gene detection, low total read length, and high mitochondrial-to-somatic gene ratio were automatically detected and filtered out using the R function “isOutlier”. The count matrix was then normalized and scaled using the “NormalizeData” and “ScaleData” functions. Principal component analysis (PCA) was performed on the top 2,000 highly variable genes using the “RunPCA” function. Significant PCA results were selected based on p-values generated by “ScoreJackStraw” to reduce background noise. Cells were clustered using the “FindClusters” function and projected onto two-dimensional space using “RunTSNE” or “RunUMAP” functions.Marker genes for each cell subtype were identified using Seurat’s FindAllMarkers function (min.pct = 0.25, logfc.threshold = 0.25). Enrichment of selected genes in these cell subtypes was studied using hypergeometric distribution tests. P-values were calculated using the phyper function (stats package version 4.2.2). In this context, N represents the number of genes expressed in the cell subtype after quality control (average expression > 0). K represents the number of expressed genes that were selected in this cell subtype. k represents the number of genes expressed in the top 200 marker genes identified for this cell subtype, and n represents the number of selected genes in the top 200 markers. The formula for calculating fold change (FC) is:

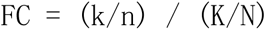

Average expression levels of each gene within each cell subtype were extracted using the AverageExpression function. The average expression matrix was then standardized using the scale function. The Heatmap function from the ComplexHeatmap v2.10.0 package was used to visualize the expression levels of marker genes across all cell subtypes. Additionally, geom_point from ggplot was used to display the selection status of the most expressed marker genes in each cell type.

### 3.10 Integrated analysis of scRNA-seq data

Prior to integrating multiple scRNA-seq datasets from two different developmental stages, we first examined for batch effects. We found that the strongest remaining factor was the difference between samples, as cells tended to cluster by sample. Therefore, we applied the R package “Harmony” (version 0.1.0)^[34]^ to correct for batch effects. Cell clusters were annotated by first identifying marker genes for each cluster using the “FindMarkers” function from the R Seurat toolkit, followed by manual annotation of cell clusters based on known cell markers for each cell type. Clusters expressing the same marker genes were merged.

### 3.11 Trajectory analysis of scRNA-seq data

To infer developmental trajectories of specific cell lineages, we performed trajectory analysis using the R package “Monocle2” (version 2.20.0) based on both scRNA-seq and scATAC-seq data. Specifically, genes differentially expressed between distinct cell types were selected for the analysis. The “DDRTree” method was then applied for dimensionality reduction and cells were ordered along pseudotime. As Monocle2 cannot a priori determine which trajectory in the tree should be designated as the “beginning”, we used the “root_state” parameter to specify the zygote stage as the starting point, followed by reapplication of “orderCells”.

### 3.12 Single-cell regulatory network analysis

To infer the regulatory activity of TFs in subpopulations, we performed gene regulatory network analysis using the pySCENIC package (version 0.11.2)^[35]^. Briefly, regulatory modules were identified by inferring co-expression between TFs and genes containing binding motifs for these TFs in their promoter regions. We generated two gene-motif ranking files, including regions within 10 kb around TSS and 500 bp upstream. The “GRNBoost” function implemented in pySCENIC was used to identify co-expression modules and quantify weights between TFs and target genes. Target genes with low positive correlation (<0.03) in each TF module were excluded from further analysis. Regulatory activity of TFs was quantified by the area under the recovery curve value for each regulon enrichment. Subpopulation-specific regulons were identified based on the average regulon activity scores within subpopulations.

### 3.13 Cell-cell communication analysis

We used the CellChat package (version 1.1.3)^[36]^ to study potential cell-cell communications and quantify intercellular communication networks. Due to incomplete annotation of receptor-ligand pairs in the bovine genome, we selected only homologous human genes for further analysis. Briefly, CellChat objects were first created using the “createCellChat” function. Communication probabilities were then inferred using the “computeCommunProb” and “computeCommunProbPathway” functions. Network centrality scores and the contribution of each ligand-receptor pair to signaling pathways were calculated using “netAnalysis_computeCentrality” and “netAnalysis_contribution”, respectively. All visualizations of cell communication networks were generated using functions implemented in the CellChat package.

### 3.14 Construction of multi-omics regulatory networks in Bactrian camels

We merged ATAC-seq libraries from four muscle samples, performed alignment, and removed duplicate BAM files. Footprint regions in muscle tissue were identified using the rgt-hint function from RGT (Regulatory Genomics Toolbox) V1.0.2. We then obtained a Seurat object with pseudo-temporal information from single-cell RNA sequencing data using the expression_profile function in the IReNA package V0.99.0. Potential regulatory interactions for each gene in the expression profile were inferred using the GENIE3 package V1.18.0^[37]^. Binding sites within footprint regions were identified using the fimo function in meme software V4.11.2. Finally, the network_analysis function generated regulatory networks connecting transcription factors to their target genes. By selecting key genes, we obtained the most critical regulatory networks in muscle tissue.

### 3.15 Motif enrichment analysis

We performed transcription factor (TF) motif enrichment analysis on differentially accessible regions (DARs) using the findMotifsGenome function from the HOMER package (version 4.11)^[38]^. This analysis was conducted to identify both known motifs and de novo motifs.

### 3.16 Dual-luciferase reporter assay

We modified the pGL3-promoter vector (Promega, E1761) by digestion with BamHI and SalI (New England Biolabs) and insertion of adaptor sequences. Similarly, we modified the pGL3-basic vector (Promega, E1751) by digestion with MluI and BglII (New England Biolabs) and insertion of linker sequences.We selected 10 putative enhancer regions from genes under selection in Bactrian camels and 10 randomly chosen negative genomic regions for analysis. Using Bactrian camel genomic DNA as a template, we amplified these enhancer and random genomic regions, each approximately 2 kb in length. The amplification products were separated on 1% agarose gels, purified, and then cloned into the modified pGL3-promoter vector.

### 3.17 Gene Ontology enrichment analysis of tissue-specific enhancers

Cis-regulatory element positions were mapped to the human genome (hg19) using the UCSC LiftOver tool with a minimum match parameter (minMatch) of 0.1 and default settings for all other parameters. Gene Ontology (GO) enrichment analysis for Biological Processes was performed on these converted regions using the Genomic Regions Enrichment of Annotations Tool (GREAT, version 3.0.0). We applied GREAT’s default association rule settings to link genomic regions to genes. Significantly enriched GO terms were identified using both the binomial test over genomic regions and the hypergeometric test over genes, with an FDR q-value threshold of 0.05.

### 3.18 Motif analysis of tissue-specific enhancer regions

We merged H3K27ac narrow peaks for each tissue using BEDTools (v2.26.097)^[39]^. Motif discovery was performed using HOMER’s findMotifsGenome.pl tool (v4.10.3). The enhancer matrix was processed with normalized read coverage changes (without quantile normalization), calculated as the average across all Bactrian camels within the same tissue, following the method described in “Identification of tissue-specific enhancers”.For each tissue, we ranked the top 5,000 enhancer H3K27ac ChIP narrow peaks by P-value and performed motif discovery on the top 3,000 peaks using HOMER’s findMotifsGenome.pl tool. Motifs with a P-value < 10^-10^ in at least one tissue and expressed in that tissue (TPM > 1) were considered significantly enriched. Similar motifs were merged using the TOMTOM tool with a P-value threshold of 10^-5^.

## 4 RESULTS

### 4.1 Genetic diversity and differentiation mechanisms in population structure

We performed deep sequencing analysis on 146 Bactrian camels from 14 breeds (2.645 Tb of data, average sequencing depth 37.74X) to investigate potential positive selection signals in the Sunite Bactrian camel genome (Fig. 1a). Alignment to the wild Bactrian camel reference genome identified approximately 15.9 million single nucleotide polymorphisms (SNPs) and 2.85 million insertion/deletion (Indel) events across the genome.

**Figure 1.**
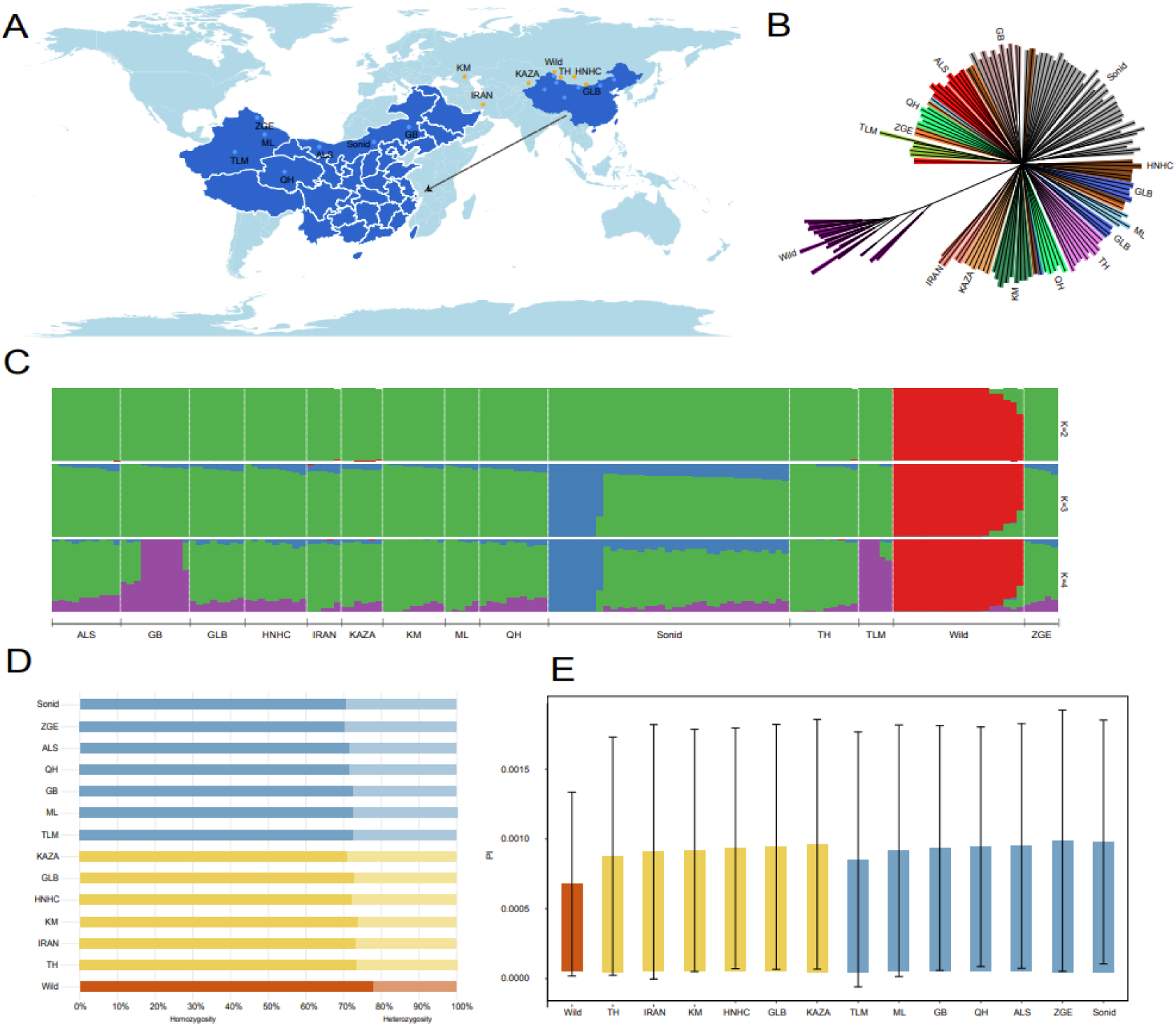
Population genetic structure of Bactrian camels based on autosomal SNP analysis. a, Geographic distribution of sampled camel populations.saralab, Phylogenetic tree with different populations represented by distinct colors.saralac, Admixture analysis showing the proportion of ancestral components for different values of K. Each color represents a distinct ancestral component, and each vertical bar represents an individual.saralad, Heterozygosity differences among populations. Blue: Chinese domestic Bactrian camels; Yellow: Foreign domestic Bactrian camels; Red: Wild Bactrian camels. Dark shades represent genomic homozygosity, while light shades indicate heterozygosity.saralae, Nucleotide diversity (π) differences among populations. Blue: Chinese domestic Bactrian camels; Yellow: Foreign domestic Bactrian camels; Red: Wild Bactrian camels.

Population structure analysis using principal component analysis (PCA) revealed clear separation between wild and all domestic Bactrian camels along the first two principal components. Some Sunite Bactrian camels also showed significant differentiation from other domestic breeds. The third principal component further distinguished Xinjiang Tarim camels from other domestic breeds (Fig. 1a, Extended Data Fig. 1). Overall, domestic camels from East, Central, West, and North Asia showed varying degrees of genetic admixture.

Neighbor-Joining (NJ) phylogenetic reconstruction confirmed the distinct branching of wild and domestic camels. Domestic breeds clustered closely, indicating close genetic relationships. Most Chinese domestic breeds formed a tight cluster, while Qinghai, Xinjiang Junggar, and Xinjiang Mulei camels showed closer genetic affinity to Mongolian domestic camels.

Admixture analysis^[40]^ revealed two ancestral components (K=2) with the lowest cross-validation error, corresponding to wild and domestic camels (Fig. 1c, Extended Data Fig. 2). This finding aligns with the two main branches in the phylogenetic tree. Notably, at least 5 wild Bactrian camel individuals showed 8.10%-35.61% domestic ancestry, possibly reflecting genetic diversity within the wild population or historical hybridization events. Conversely, small proportions (0.22%-1.06%) of wild Bactrian camel genetic components were detected in Central Asian domestic populations, including Inner Mongolia Alxa (ALS), Iran Ardabili (IRAN), Kazakhstan Almaty (KAZA), and Mongolian Tokhom-tungalag (TH).

Further population differentiation analysis revealed distinct genetic components in at least 7 individuals from the Sunite (Sonid) Bactrian camel population (K=3). Similarly, unique genetic signatures were observed in the Inner Mongolia Gobi Red (GB) and Xinjiang Tarim (TLM) camel populations (K=4).

To investigate potential migration events between populations, we employed the TreeMix[41] model. We tested migration events (m) ranging from 1 to 10 and determined m=4 as the optimal number based on the rate of change in the second-order derivative of the log-likelihood values. This model explained 98.23% of the genetic variation (Extended Data Fig. 3). The maximum likelihood tree suggested four possible migration paths: from Sunite camels to Iranian Ardabili and Xinjiang Junggar camels, from Xinjiang Mulei to Xinjiang Junggar camels, and from wild Bactrian camels to Kazakhstan Almaty camels. However, the residual heatmap did not indicate strong migration signals, suggesting only a degree of genetic admixture between these camel populations without clear migration footprints (Extended Data Fig. 4 and 5).

To compare genetic diversity among different camel populations, we first assessed their genomic heterozygosity (Fig. 1D). Our analysis revealed that domestic camel populations in China exhibited slightly higher genomic heterozygosity than their foreign counterparts (average heterozygosity of 0.28 and 0.27, respectively), although this difference was not statistically significant. In contrast, wild camels showed a markedly reduced genomic heterozygosity, with an average of approximately 0.22, lower than that of domestic camels.

Furthermore, we calculated nucleotide diversity (π) values (Fig. 1E) and found that Chinese domestic camels displayed marginally higher nucleotide diversity compared to international domestic populations (π = 0.89 × 10^-3^ and 0.87 × 10^-3^, respectively). Wild camels, however, exhibited significantly lower nucleotide diversity than domestic camels, with an average π value of 0.63 × 10^-3^, consistent with the genomic heterozygosity estimates.

### 4.2 Deep annotation and mechanistic analysis of population genomic variations

Theoretical predictions suggest that wild populations should exhibit higher genetic diversity than their domestic counterparts. However, elevated genetic drift or inbreeding in small wild populations may lead to reduced genetic diversity compared to domestic groups. We investigated this phenomenon in camel populations using F-statistics (Fst) based on the method proposed by Weir and Cockerham^[42]^.Our analysis revealed substantial genetic differentiation between wild camels and all domestic populations, with Fst values ranging from 0.21 to 0.29 (Fig. 2a). Among domestic camels, Tarim (TLM) and Iranian (IRAN) populations showed moderate levels of genetic differentiation from other domestic groups, with average Fst values of approximately 0.07 and 0.06, respectively. Overall, genetic differentiation among domestic camel populations ranged from negligible to moderate (Fst values ∼0-0.09).

**Figure 2.**
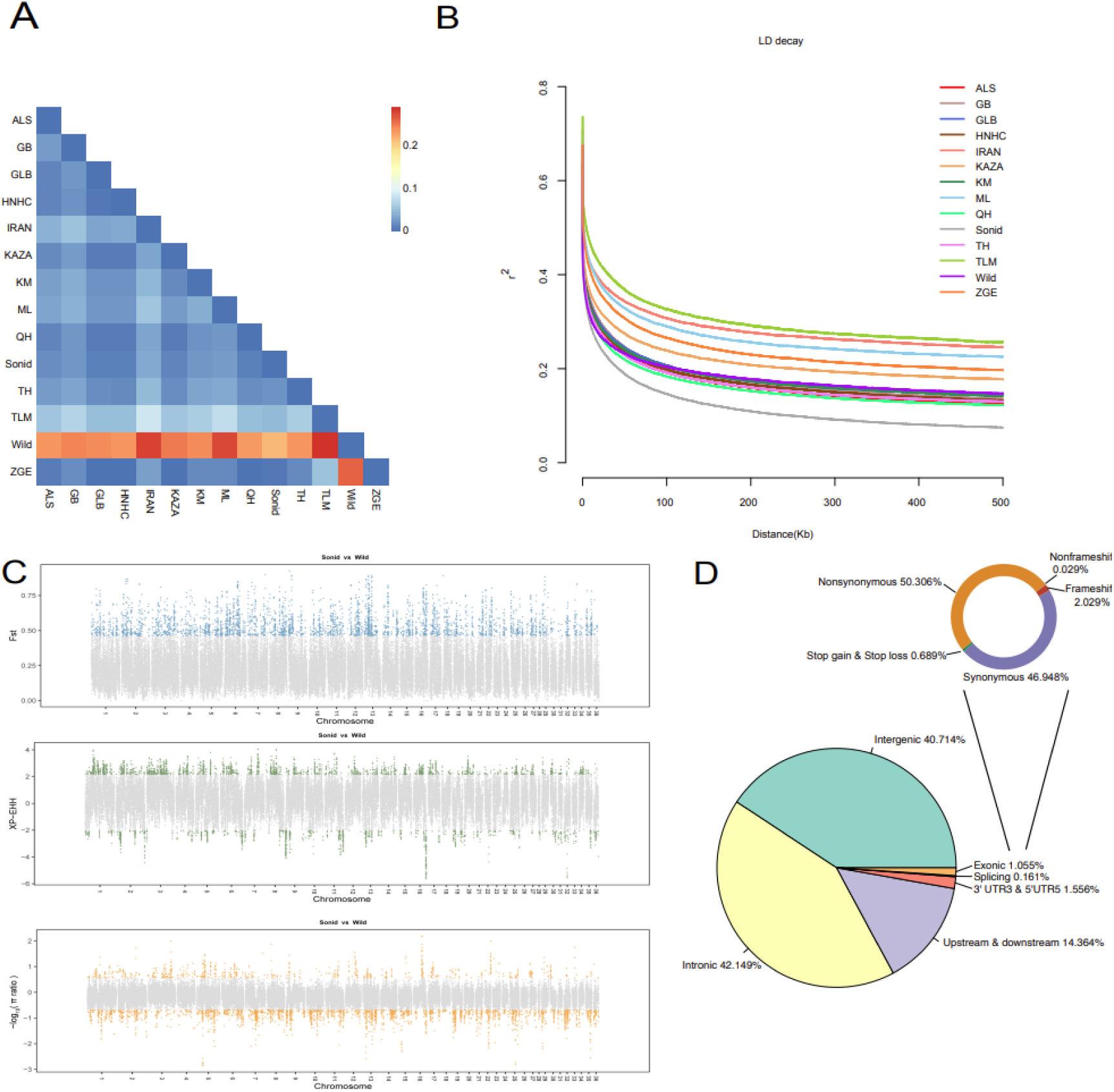
Comprehensive analysis of camel population genetic structure and natural selection based on statistical methods. a, Pairwise population Fst. The heatmap represents average Fst values calculated using 50 kb windows with 25 kb steps. Wild camels show substantial genetic differentiation compared to all domestic populations.saralab, Linkage disequilibrium (LD) decay. Each color represents a distinct population.saralac, Identification of selected regions using three statistically robust methods: population differentiation index (F-statistics, Fst), nucleotide diversity (π), and cross-population extended haplotype homozygosity (XP-EHH).saralad, Functional annotation of variant sites in Sunite camels. The distribution of these annotations across the genome reveals that the majority of variant sites are located in non-coding regions.

We investigated linkage disequilibrium (LD) patterns across camel populations by calculating the LD coefficient (r^2) between pairs of SNPs within 500 kb windows and plotting LD decay curves (Fig. 2b). All camel populations exhibited similar overall trends in LD decay, with r^^2^ values gradually decreasing as the physical distance between SNP pairs increased.

However, significant variations in the rate and extent of LD decay were observed among populations, reflecting their diverse genetic backgrounds. The Sonid Bactrian camels showed the most rapid LD decay (approximately 2.20 kb), while the Tarim camels exhibited the slowest decay (approximately 50.00 kb). These decay rates are influenced by multiple factors, including recombination rates, natural selection, genetic drift, and mutation rates.

Domestication has led to significant behavioural and phenotypic differences between wild and domestic camel populations, driven by human-mediated selection for economically important traits. To identify genomic regions under positive selection in Sonid camels, we employed three widely-recognized statistical methods: F-statistics (Fst), nucleotide diversity (π), and cross-population extended haplotype homozygosity (XPEHH).

Our analysis revealed 3,821, 742, and 2,458 potentially selected regions in Sonid camels compared to wild camels using Fst, π, and XPEHH, respectively (Fig. 2c). To enhance the robustness of our findings, we focused on 1,590 regions identified by at least two methods, which were annotated to 423 candidate genes.

KEGG pathway enrichment analysis of these candidate genes (Extended Data Fig. 3b) highlighted their involvement in adipocytokine signalling (including PRKAB2, G6PC, PPARGC1A, G6PC3, STAT3) and prolactin signalling pathways (including STAT5B, MAPK1, STAT3, SOS1). These pathways are likely associated with key economic traits in Sonid camels.

Conventionally, annotation of coding variants aids in interpreting selection signatures. However, our analysis revealed that 98.85% of annotated SNPs and indels were located outside exonic regions. Strikingly, within the putatively selected regions, 99.94% of variants were similarly positioned outside exons. This observation underscores the critical role of non-coding regions in camel genetics, suggesting their significant influence on trait selection and providing crucial insights for comprehensive understanding of camel genome function and genetic improvement.

### 4.3 Overview of epigenetic data

The majority of genetic variants reside in non-coding regions, presenting significant challenges for functional annotation and interpretation of their role in natural selection. To elucidate how non-coding variants influence phenotypic differences between wild and Sonid camels, we conducted an integrative analysis of gene expression and chromatin accessibility across multiple tissues (Fig. 3a).

**Figure 3.**
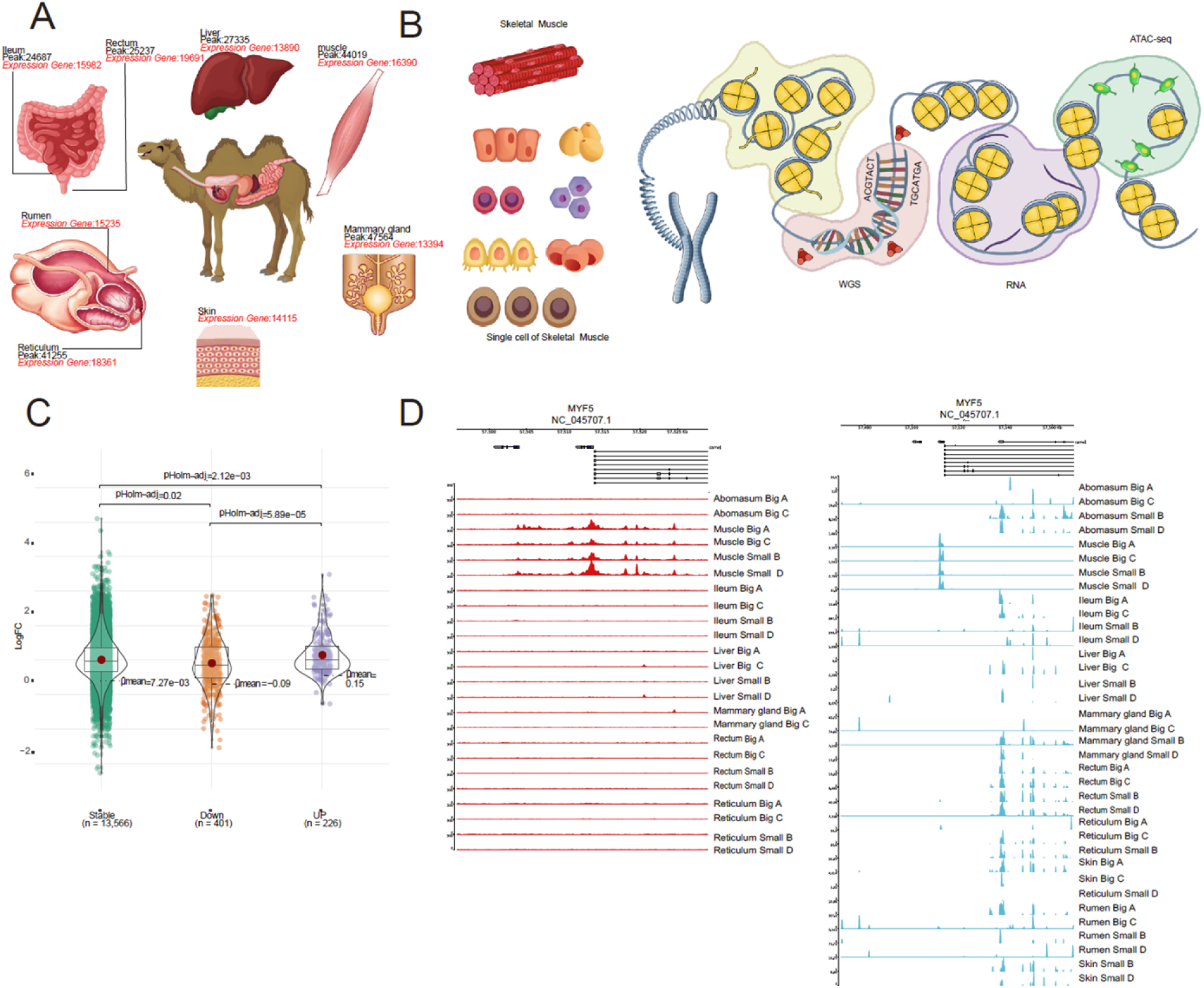
Gene expression and regulatory mechanisms in Sonid camels. a, Experimental design and sample collection for RNA sequencing (RNA-seq) and Assay for Transposase-Accessible Chromatin using sequencing (ATAC-seq) analyses in Sonid camels.saralab, Identification of expressed genes and regulatory regions (peaks) through integration of whole-genome sequencing (WGS), RNA-seq, and ATAC-seq data, providing a comprehensive foundation for functional studies.saralac, Changes in enhancer activity associated with differentially expressed genes. The fold change in signal intensity is shown, with ** indicating statistical significance (p < 0.01), revealing key alterations in enhancer activity during expression regulation.saralad, Expression and regulation of the MYF5 gene. ATAC-seq (red) and RNA-seq (blue) profiles of the MYF5 promoter and gene body regions, illustrating the activity patterns in transcriptional regulation.

We performed RNA sequencing (RNA-seq) on eight tissues (rumen, reticulum, rectum, ileum, mammary gland, skin, muscle, and udder) from Bactrian camels. Additionally, we carried out Assay for Transposase-Accessible Chromatin using sequencing (ATAC-seq) on six adult and five juvenile camel tissues (reticulum, rectum, ileum, mammary gland, skin, and muscle), with two biological replicates per tissue. In total, we analyzed 94 sequencing datasets from major tissues of adult and juvenile camels (Fig. 3a).

This comprehensive approach generated approximately 3.968 billion raw sequencing reads, of which 3.435 billion were successfully mapped to the reference genome. After alignment and filtering, we retained an average of 75.36% of the reads (Supplementary Table 2, ATAC). Further analysis revealed an average of 35,213 open chromatin peaks and 16,803 detectable genes (TPM ≥ 1) across all tissues for ATAC and RNA-seq, respectively. On average, open chromatin regions accounted for 3.86% of the entire genome in each library (Fig. 3b).

Using ATAC-seq, we identified candidate enhancers and promoters across various tissues, with an average of 4,012 enhancers and 10,853 promoters per tissue. Enhancer sequences play a crucial role in shaping tissue-specific gene expression patterns. We summarized tissue-specific enhancer patterns in adult and juvenile camel tissues (Supplementary Fig. 4a) and identified 30,582 tissue-specific enhancers. A heatmap of the enhancer signal matrix revealed distinct tissue-specific intensity patterns (Supplementary Fig. 4d).

HOMER motif analysis demonstrated that these tissue-specific enhancers and promoters were enriched for key motifs closely associated with the development of their respective tissues. For instance, longissimus dorsi muscle-specific enhancers were enriched for muscle-specific transcription factors such as Mef2a and Mef2b, which have been shown to influence myotube formation43 and play important roles in skeletal muscle regeneration44.

Gene Ontology (GO) analysis of genes adjacent to tissue-specific regulatory elements revealed functional enrichment closely related to the corresponding tissue processes, such as muscle differentiation in muscle tissue and response to nutrient levels in the rectum (Supplementary Fig. 4e).

These findings provide important insights into tissue-specific regulatory mechanisms and further confirm the role of these identified sequences as tissue-specific cis-regulatory elements.

We investigated the transcriptomic features of eight tissues from adult and juvenile Bactrian camels using RNA sequencing (RNA-seq). The RNA expression patterns in these tissues were diverse and clustered into 17 groups using the kmeans clustering function in R (Supplementary Fig. 5a). A heatmap of expression patterns revealed distinct tissue specificity, with over half of the clusters showing significant tissue-specific trends. Notable differences in expression levels were also observed between adult and juvenile camels in the same tissues (Supplementary Fig. 5c).

Further analysis of the normalized gene expression matrix identified an average of 826 tissue-specific genes in each tissue of adult and juvenile camels, with significant expression differences between adult and juvenile tissues. Gene Ontology (GO) enrichment analysis revealed significant enrichment of these tissue-specific genes in various tissue-specific functions (Supplementary Fig. 5b). For example, specific genes in adult camel skin tissue were primarily enriched in biological processes such as skin epidermis development, hair follicle development, and hair cycle. In contrast, specific genes in the longissimus dorsi muscle tissue were significantly enriched in processes such as myocyte differentiation.

We found a close association between tissue-specific enhancers and gene-specific expression. Integrated analysis of RNA-seq and ATAC-seq data demonstrated clear tissue specificity and synergy between candidate enhancer signals and transcriptomic signals. For instance, the MyF5 gene, which plays multiple roles throughout muscle development, from early muscle progenitor cell determination to myotome formation, and adult muscle maintenance and regeneration, potentially influencing final muscle traits^[45–47]^, showed open chromatin regions only near this gene in the longissimus dorsi muscle tissue, with weak ATAC-seq signals near MyF5 in other tissues. Correspondingly, this gene also displayed significant tissue-specific signals in the RNA-seq data, indicating active transcription in the longissimus dorsi muscle tissue (Fig. 3d).

These findings strongly support the critical role of tissue-specific genes in different biological processes and highlight the relationship between tissue-specific enhancers and gene expression.

Further analysis of differentially expressed genes (DEGs) between adult and juvenile camels revealed the most significant number of DEGs in the mammary gland, with 3,163 upregulated and 4,168 downregulated genes. The functional characteristics of DEGs in each tissue were closely related to the phenotypes of the two camel breeds.

For instance, in rectal tissue, 901 genes were upregulated and 1,348 genes were downregulated in adult camels compared to juvenile camels (Supplementary Fig. 3b). Upregulated genes were significantly enriched in biological processes such as digestive tract development, transmembrane transport, and actin filament-based movement in reticulum function. Downregulated genes were significantly enriched in immune cell differentiation processes (see Supplementary Table: Differential Gene Table).

Based on these observations, we hypothesized that changes in regulatory element activity might be a key factor driving differential gene expression between adult and juvenile camels. We calculated the fold change of enhancer signals near differentially expressed genes in adult and juvenile camels. Taking the longissimus dorsi muscle tissue as an example, Welch’s test results showed that the fold change of enhancer expression near upregulated genes was significantly higher compared to downregulated or stable genes (p.adj = 5.89e-05; p.adj = 2.12e-03). This finding emphasizes the crucial role of enhancer activity changes in gene expression differences during camel tissue development (Fig. 3c). Our generated transcriptomic and epigenomic data accurately captured this process, providing valuable insights into the molecular mechanisms underlying tissue-specific gene regulation and developmental changes in Bactrian camels. These findings contribute to our understanding of the genetic basis for adaptive traits in these animals and may have implications for improving camel breeding and management strategies.

Finally, in addition to investigating tissue-specific gene expression, we aimed to explore gene co-expression patterns and expression stability across tissues. We employed weighted gene co-expression network analysis (WGCNA) to identify gene modules associated with specific traits. We successfully identified modules significantly correlated (positively or negatively) with different tissues and extracted module genes positively correlated with skin, mammary gland, and longissimus dorsi muscle for subsequent Gene Ontology (GO) enrichment analysis.

Specifically, the orange-red module (correlation = 0.48, P < 0.01) and red module (correlation = 0.83, P = 1 × 10^-9^), associated with skin phenotypes, were enriched for critical biological processes including skin epidermis development, epidermal development, and body fluid level regulation. The magenta module (correlation = 0.97, P = 3 × 10^-21^), closely related to longissimus dorsi muscle, was enriched for pathways involving muscle cell differentiation, striated muscle contraction, muscle cell development, and muscle organ development. The dark grey module (correlation = 0.71, P = 3 × 10^-6^), associated with mammary gland phenotypes, was enriched for pathways such as positive regulation of lipid localization. These analyses revealed that gene co-expression studies can provide deeper insights into trait-related gene sets.

Furthermore, we investigated the enrichment of genes under selection at the co-expression level. Hypergeometric distribution tests revealed significant enrichment of genes under selection (224 genes) in the cyan module (P = 1.93 × 10^-2^) and turquoise module (P = 7.63 × 10^-10^). Notably, these modules showed weak positive correlations (correlation ≤ 0.4) with multiple traits, suggesting that some genes under selection may exhibit consistent expression patterns across different tissues.

WGCNA revealed potential interaction effects between genes in specific tissues, such as the promoting effect of upstream elements at transcription factor binding sites. We believe that further mining of cross-tissue, multi-omics data will contribute to constructing a more comprehensive gene regulatory network.

### 4.4 Integration of genetic selection signals in epigenomic analyses

To explore the expression patterns of genes under selection, we constructed an expression heatmap of selected genes. Our results revealed distinct tissue-specific expression patterns for most selected genes (Fig. 4A). For example, SOX6 and RIMKLA showed specific expression in the longissimus dorsi muscle due to selection effects. SOX6, a key transcription factor, plays a crucial role in muscle development, fiber type determination, and transformation^[48]^. RIMKLA encodes another N-acetylaspartylglutamate (NAAG) synthetase, potentially influencing muscle activity indirectly through neuronal function [49]. Some selected genes also exhibited high expression levels across multiple tissues, consistent with our module analysis results.

**Figure 4.**
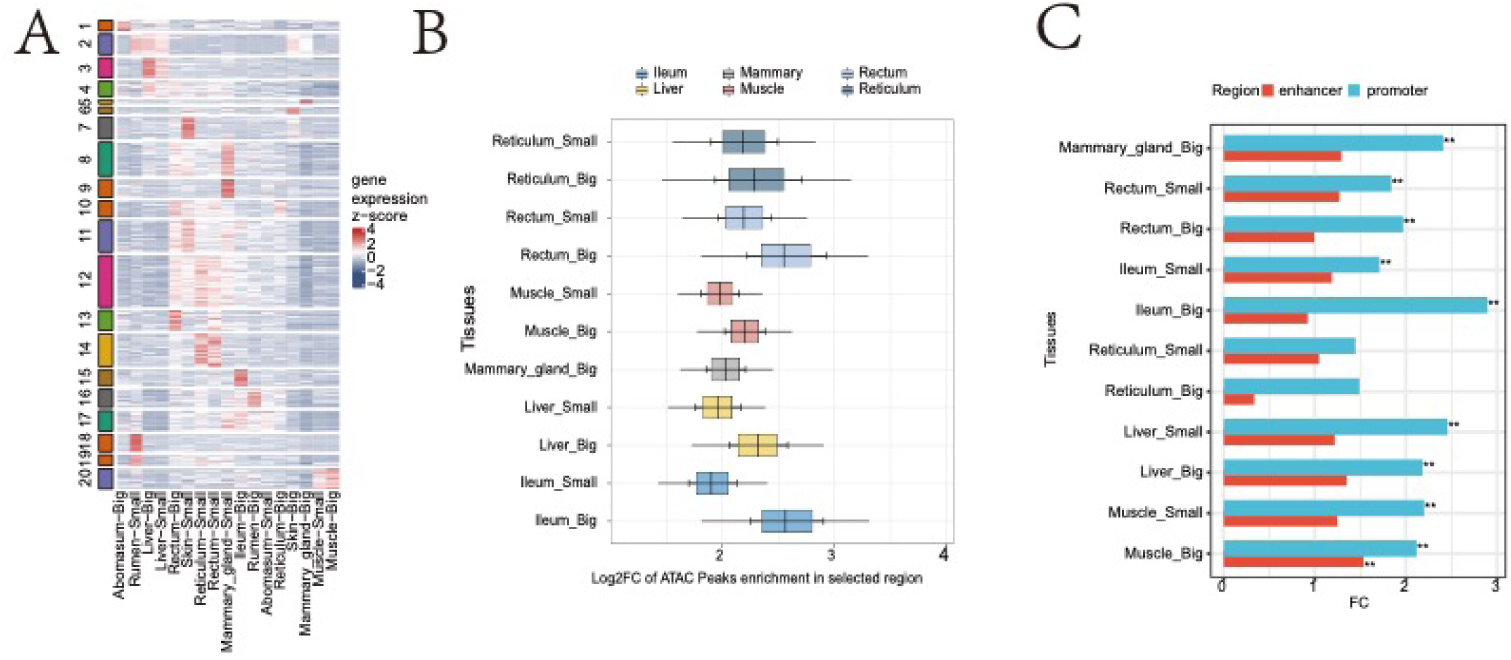
Expression patterns and regulatory region enrichment analysis of genes under selection in Sunite camels. A) Expression profile of genes under selection across various tissues in Sunite camels: This heatmap reveals expression differences of specific genes across different tissues, likely resulting from natural selection pressures.saralaB) Significant enrichment of open chromatin regions in genes under selection: This panel demonstrates the enrichment of open chromatin regions within genes under selection in Sunite camels. This suggests these regions may play a crucial role in regulating gene expression.saralaC) Enrichment analysis of enhancers and promoters in natural selection signals: This bar plot shows the enrichment of enhancer and promoter regions in genes affected by natural selection. ** indicates p < 0.01, * indicates p < 0.05, reflecting the critical role of these regulatory elements in genetic adaptation.

**Figure 5:**
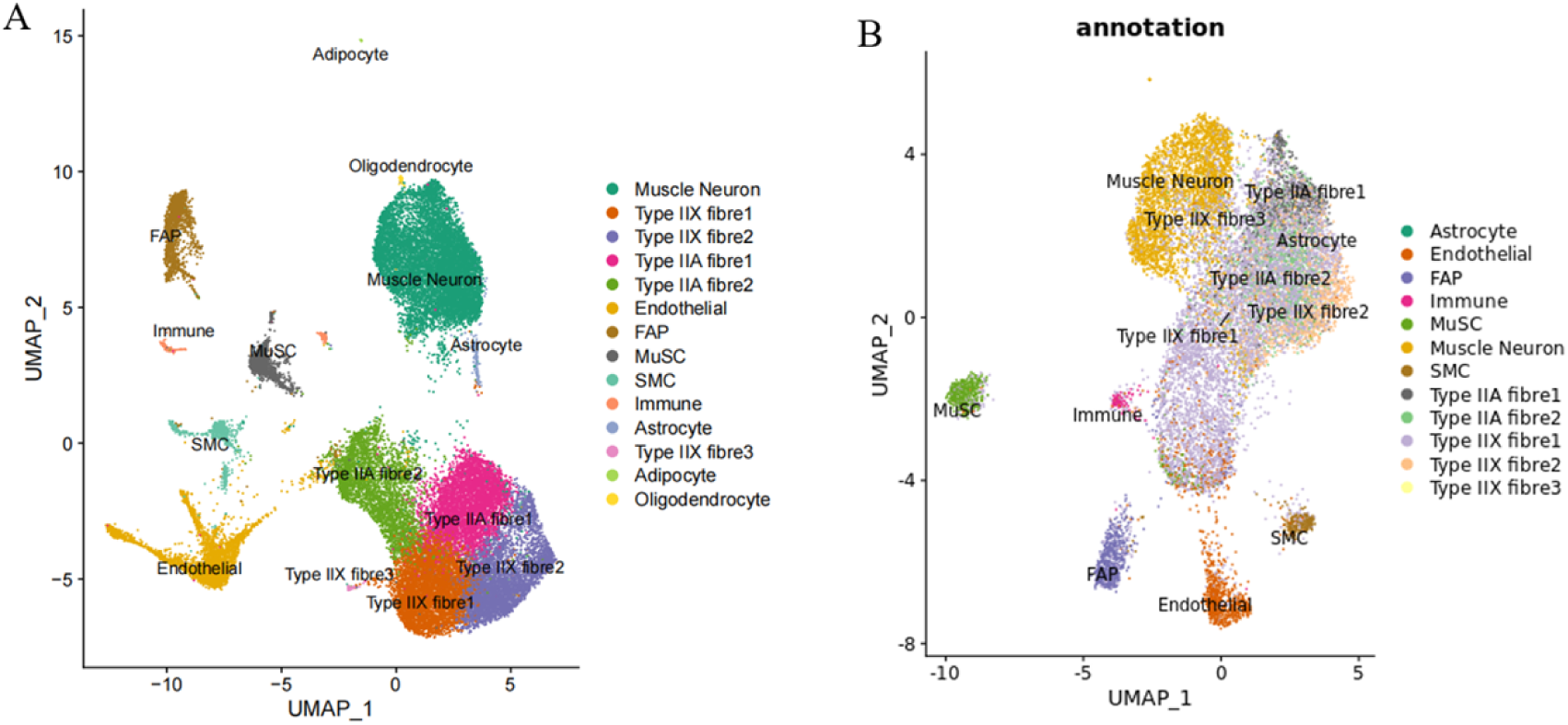
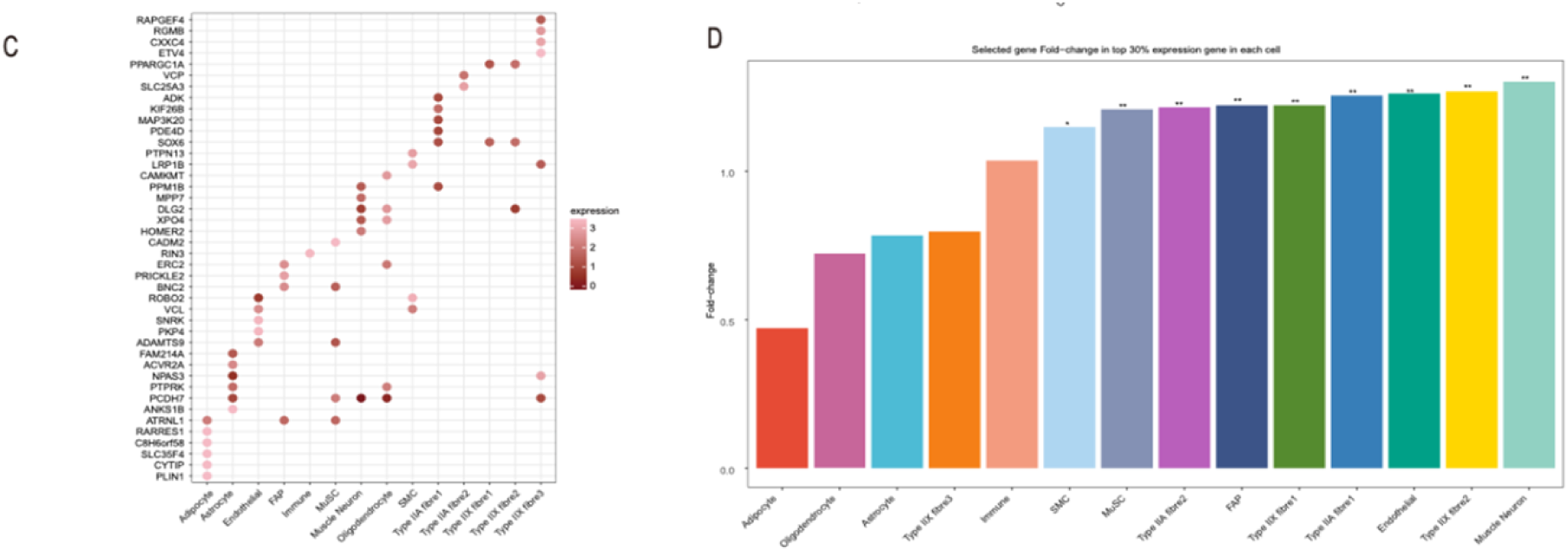
Single-cell analysis reveals cellular heterogeneity and activity changes in selected genes and regulatory elements. a, Single-cell RNA-seq visualization. Cells are color-coded by tissue origin, demonstrating cellular diversity and tissue specificity.saralab, Single-cell ATAC-seq visualization. Cells are color-coded by tissue origin, illustrating cellular diversity and tissue specificity.saralac, Analysis of selected marker genes. Identification of marker genes under evolutionary or functional selection pressure.saralad, Selective enrichment of cell type-specific gene expression. Displays the selective enrichment of genes with expression levels in the top 30% for each cell type at the cellular level. Double asterisks indicate P < 0.01; single asterisk indicates P

Given that most SNPs and indels in selected regions were located in non-coding areas, we hypothesized that these regions might be enriched in identified open chromatin regions. To test this hypothesis and exclude the influence of highly variable functional coding regions, we calculated enrichment indices for selected regions in open chromatin areas across different tissues. Further analysis revealed significant enrichment of open chromatin regions from all tissues within selected areas (Fig. 4B, Log2FC>1, P<0.01).

This finding led us to speculate that cis-regulatory elements might also be enriched in selected regions. To understand the potential roles of proximal and distal cis-regulatory elements in natural selection, we performed permutation tests on promoters and enhancers to calculate enrichment indices. Results showed significant enrichment of promoters from multiple tissues and both promoters and enhancers from adult camel longissimus dorsi muscle in selected regions, suggesting that cis-regulatory elements associated with important economic traits experienced strong selection pressure during Sunite camel domestication (FC>1, P<0.01) (Fig. 4C).

This enrichment may result from strict artificial selection for meat-related economic traits, such as meat yield and quality, specifically targeting cis-regulatory elements associated with skeletal muscle development. Our findings emphasize the critical role of cis-regulatory elements in domestication and adaptive evolution, revealing their profound impact on camel skeletal muscle growth and development regulation.

### 4.5 Integrative analysis of genetic selection signals in single-cell genomics

To elucidate the genetic regulatory mechanisms by which selected genes influence muscle in Sunite camels, we focused on a detailed study of skeletal muscle cell subpopulations. Our aim was to reveal the expression patterns of selected genes across different cell subsets for a more precise understanding of their regulatory mechanisms.

We analysed the longissimus dorsi muscle tissue from two adult and two juvenile camels using single-cell RNA sequencing (scRNA-seq). After integrating all single-cell sequencing samples and removing batch effects using integration functions, we performed dimensionality reduction and unsupervised clustering on 31,961 cells, ultimately identifying 19 distinct cell clusters. Following differential expression analysis and examination of selected marker genes (Top 100, average log2 fold change ≥1, adjusted P value ≤0.01), we manually annotated 14 different subpopulations, including Type IIA fibres, Type IIX fibres, endothelial cells, muscle neurons, fibro-adipogenic progenitors (FAPs), muscle stem cells (MuSCs), smooth muscle cells (SMCs), immune cells, astrocytes, adipocytes and oligodendrocytes. A heatmap of marker gene expression further validated the accuracy of these cell annotations.

To investigate regulatory events during myocyte development, we analysed single-cell chromatin accessibility profiles. Using a clustering algorithm based on shared nearest neighbour (SNN) modularity optimization, we identified 12 distinct differential accessibility peak clusters. We focused on marker genes affected by selected regions to explore key genes potentially regulating skeletal muscle cell composition in Sunite camels.

We found that most selected marker genes showed specificity in particular cell subsets, although some genes such as SOX6 and PCDH7 were marked as marker genes in multiple cell populations despite being selected genes, suggesting they may play important roles across multiple cell types. Additionally, by focusing on the top 30% of expressed genes in each cell subset, we calculated the enrichment index of selected genes in various cell subpopulations. Notably, clusters associated with muscle fibre cells, muscle stem cells and skeletal muscle cells showed significant enrichment of selected genes, while clusters such as immune and fat cells did not exhibit such enrichment.

Furthermore, we analysed the dynamic changes in gene expression across different cell subpopulations and the potential for these genes to recruit transcription factors near open chromatin regions, constructing a core regulatory network focused on Sunite camel skeletal muscle. This network showed biological functional enrichment in processes related to skeletal muscle growth and fatty acid synthesis, highlighting its crucial role in camel meat quality traits.

Within the network, the core transcription factor SOX6 was involved in regulating skeletal muscle growth and development processes, underscoring the immense potential of epigenetics and single-cell transcriptomics in revealing genes associated with growth and development. Simultaneously, this finding emphasizes the importance of selection effects in non-coding regions in influencing biological traits, providing new perspectives for a deeper understanding of genetic regulatory networks.

### 4.6 Integrated analysis of genetic selection signals in single-cell developmental and differentiation trajectories

Pseudotime analysis of muscle cell subpopulations in the four-humped Sunit Bactrian camel revealed differentiation trajectories originating from satellite cells and diverging towards type IIa and IIx muscle fibers (Fig. 6a). Dynamic expression patterns of candidate genes in selected regions across muscle cell subpopulations showed that ARHGAP28 is specifically highly expressed in muscle stem cells but not in other subpopulations (Fig. 6c). This may reflect its unique role in maintaining stem cell properties, regulating muscle tissue development and regeneration, and determining cell fate. ARHGAP28 is highly expressed in the early stages of pseudotime development but not expressed at the end of development (Fig. 6d).

**Figure 6.**
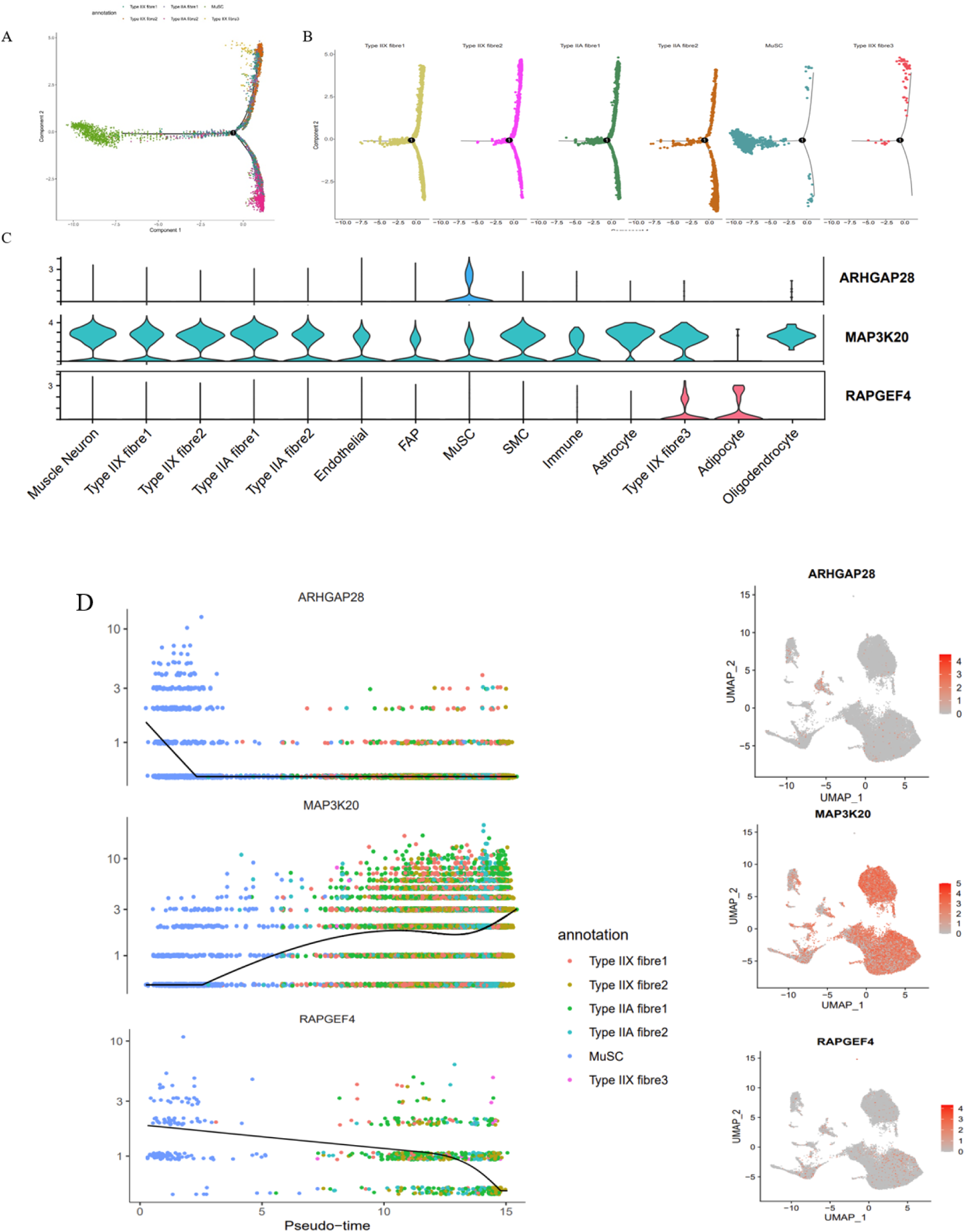
Reconstruction of myogenic differentiation trajectory using pseudotime analysis. a, Pseudotime analysis of myogenic cells using Monocle 2.saralab, Visualization of myogenic differentiation trajectory colored by cell identity.saralac, Violin plots showing the expression of candidate regulatory genes near selected mutation sites in each cell cluster. ARHGAP28, MAP3K20, and RAPGEF4 are important candidate genes affecting muscle development.saralad, Expression patterns of candidate regulatory genes across different cell clusters.

Studies have shown that MAP3K20 mediates the MAP kinase signaling pathway induced by skeletal muscle contraction^[50]^. Although MAP3K20 may not be highly expressed in intramuscular fat cells, its high expression in other cell types may be associated with key processes such as cellular stress response, differentiation, proliferation, and survival. For example, it may be involved in neuroprotection in neurons and glial cells; in muscle fibers, it may regulate muscle contraction and metabolism; in endothelial cells and smooth muscle cells, it may affect vascular function; in progenitor cells such as fibro-adipogenic progenitors and muscle satellite cells, it may regulate differentiation and regeneration processes; in immune cells, it may participate in inflammatory responses. This widespread expression pattern suggests that MAP3K20 may play an important role in maintaining normal physiological functions of muscle tissue and responding to various stimuli. However, these speculations need to be verified through further experimental studies. MAP3K20 is not expressed in the early stages of development but is highly expressed in the late stages of development (Fig. 6d).

The specific high expression of RAPGEF4 in Type IIX fiber cells and adipocytes reveals its key role in muscle function and fat metabolism. Studies have shown that RAPGEF4 may affect muscle function by regulating Rap1 activity^[51]^. In Type IIX fibers, RAPGEF4 may be involved in regulating rapid muscle contraction, energy metabolism, and fiber type maintenance; while in adipocytes, it may influence fat metabolism, insulin sensitivity, cell differentiation, and inflammatory responses. This dual expression pattern suggests that RAPGEF4 plays an important role in coordinating whole-body energy balance, metabolic flexibility, and exercise performance.

The RAPGEF4 gene shows a significant early-to-mid-stage high expression pattern during muscle cell differentiation (Fig. 6d). This unique expression pattern reveals that RAPGEF4 may play distinctly different roles at different stages of myogenesis. In the stem cell stage, high expression of RAPGEF4 may be crucial for maintaining stem cell properties or triggering the initiation of differentiation. The second expression peak in the middle stage of differentiation suggests its important role in promoting muscle cell maturation or maintaining the differentiated state. This spatiotemporal-specific expression pattern provides new perspectives for understanding the multiple functions of RAPGEF4 in the process of myogenesis, while also opening up potential pathways for targeted regulation of muscle development.

To explore the mechanisms of selection effects on highly expressed genes in cell subpopulations, we hypothesized that the selection effects on muscle traits may not only originate from a few SNPs and Indels causing missense or frameshift mutations. We propose that these effects might also involve SNPs and Indels on selected elements, potentially altering the activity of cis-regulatory elements or affecting their binding affinity with transcription factors, thus creating differences in muscle traits between wild and Sunit camel populations.

To test this hypothesis, we selected genes with expression levels in the top 30% in skeletal muscle stem cells and analyzed their upstream and downstream cis-regulatory elements. We then focused on highly differentiated variant sites (delta AF>0.6) between wild and Sunit camels within these cis-regulatory elements. Combining single-cell RNA-seq and ATAC-seq data analysis, we identified MAP3K20, RAPGEF4, and MTFR1L as important candidate genes.

To further validate the impact of these mutations on element activity, we conducted luciferase reporter gene assays in C2C12 cell lines. Among the 8 pairs of enhancers tested, three pairs showed significantly higher activity in the Sunit camel mutant enhancers compared to the reference enhancers (T-test, P<0.01). Notably, two pairs of enhancers (chr5:46451711|A−G, chr13:30361170|T−C) (Fig. 7b,c) underwent a significant regulatory element identity shift from silencers to enhancers, a relatively rare phenomenon that provides strong experimental evidence for understanding how mutation sites can alter element activity and function.

**Figure 7.**
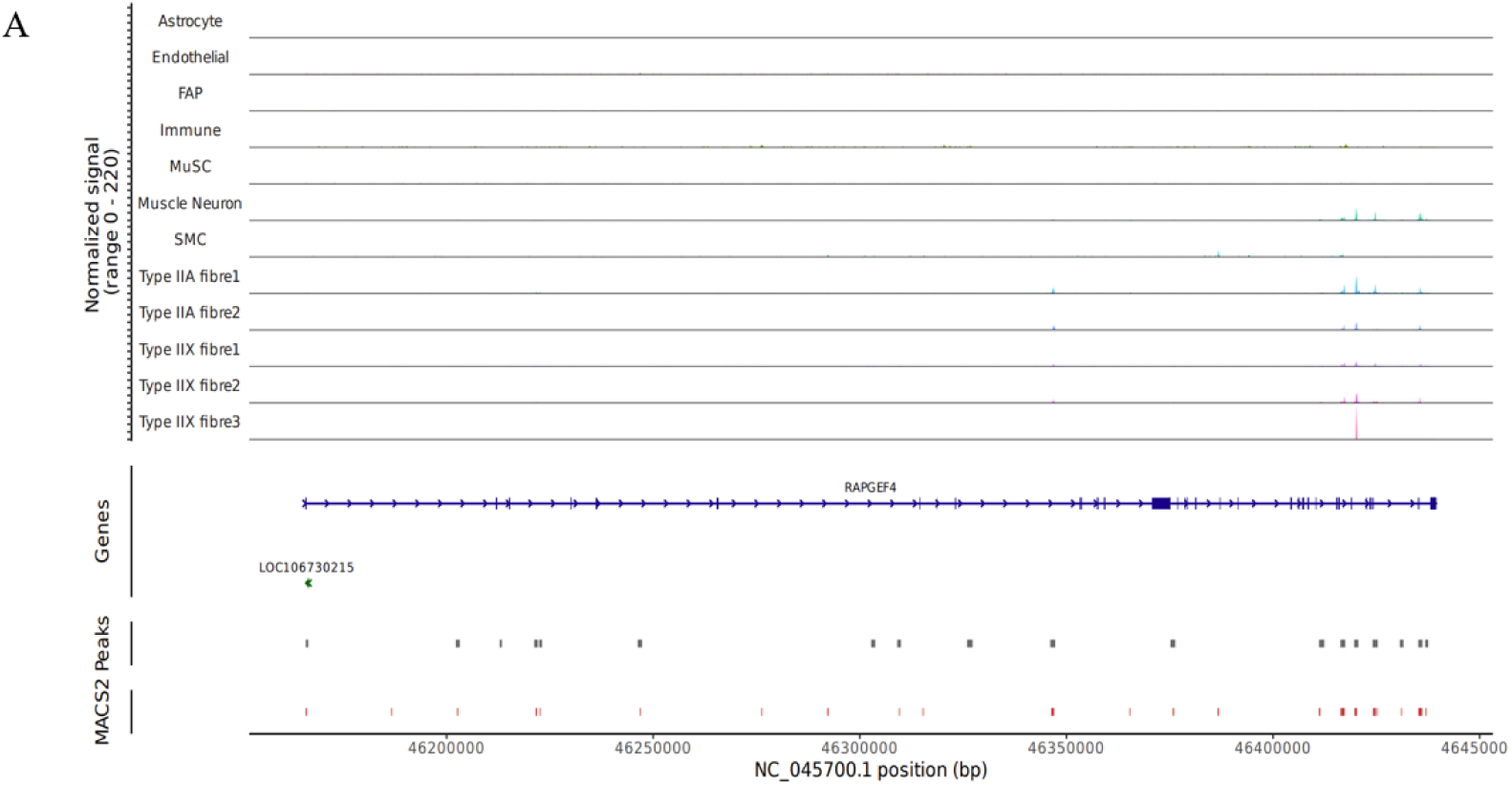
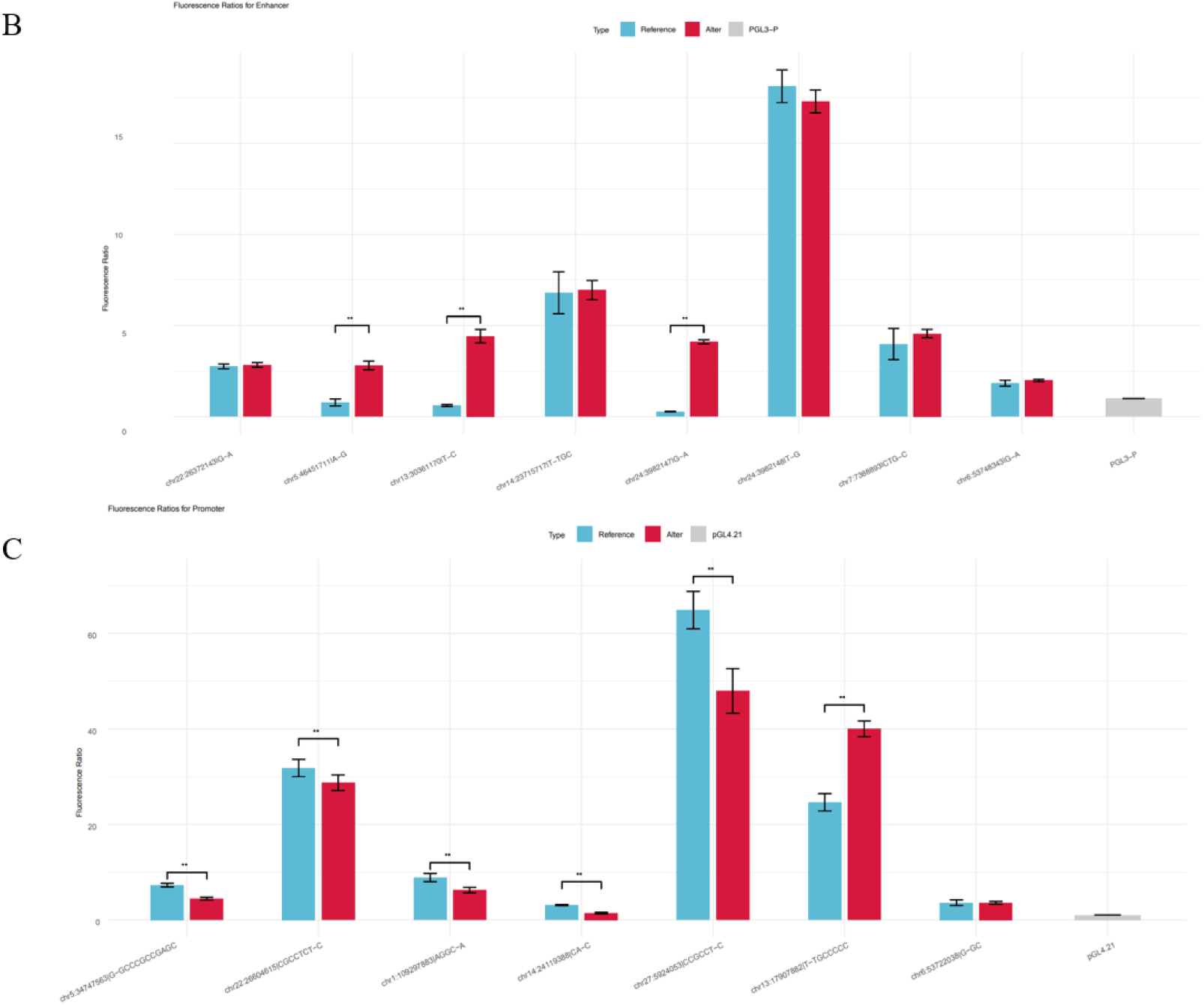
Single-cell ATAC peak analysis of RAPGEF4 gene in Bactrian camel skeletal muscle. a, Scatter plot showing the correlation between RAPGEF4 gene expression levels and ATAC-seq signal intensity in its promoter region.saralab,c, Analysis of the selective effects of SNP and Indel mutations on the activity of promoters (b) and enhancers (c). The changes in activity before and after SNP and Indel mutations under selective effects are explored.sarala** indicates p < 0.01, * indicates p < 0.05.

Among these, the candidate regulatory gene MAP3K20 near chr5:46451711|A−G is known to be closely related to muscle fiber differentiation. Studies have found that MAP3K20 participates in regulating the JNK pathway, affecting muscle development and regeneration processes[52], and plays a crucial role in contraction-induced signaling in skeletal muscle[53].

Of the 7 pairs of promoters tested, 6 pairs showed significant differences in activity between mutant and reference promoters (T-test, P<0.01). Notably, the Indel chr13:17907882|T−TGCCCCC significantly increased promoter activity post-mutation. Its candidate target gene, MTFR1L, has been reported to influence mitochondrial morphology through mediating phosphorylation[54], potentially affecting muscle performance by influencing mitochondrial distribution and function.

Future in-depth gene interference experiments will further reveal the specific regulatory mechanisms and biological significance of these genes in muscle tissue. Integrated analysis of single-cell RNA-seq and ATAC-seq data indicates that the candidate gene RAPGEF4 at chr5: 46451671|T−A plays an important regulatory role in muscle subpopulations (Fig. 7a).

We selected the muscle differentiation and development-related gene RAPGEF4 for tissue localization and expression verification. Immunohistochemical analysis revealed that RAPGEF4 is primarily localized in the cell membrane of the longissimus dorsi muscle tissue in Bactrian camels, with expression also observed in the cytoplasm (Fig. 8a). To further investigate the role of RAPGEF4 in muscle differentiation, we constructed RAPGEF4 overexpression (OE-RAPGEF4) and interference (si-RAPGEF4) lentiviral vectors in C2C12 cells, using an empty vector as a control. We measured the expression levels of myogenic marker genes MYOD and MYOG by quantitative PCR on days 0, 3, 5, and 7 of myogenic induction (Fig. 8b). The results showed that as the induction time increased, the expression levels of MYOD and MYOG in the OE-RAPGEF4 group were significantly higher than those in the control group (P < 0.01, two-tailed t-test, n = 3 biological replicates). Conversely, the expression levels of these two genes in the si-RAPGEF4 group were significantly lower than those in the control group (P < 0.01, two-tailed t-test, n = 3 biological replicates). These findings suggest that RAPGEF4 expression levels positively correlate with the degree of myogenic differentiation, indicating that RAPGEF4 may play an important role in promoting muscle differentiation.

**Figure 8.**
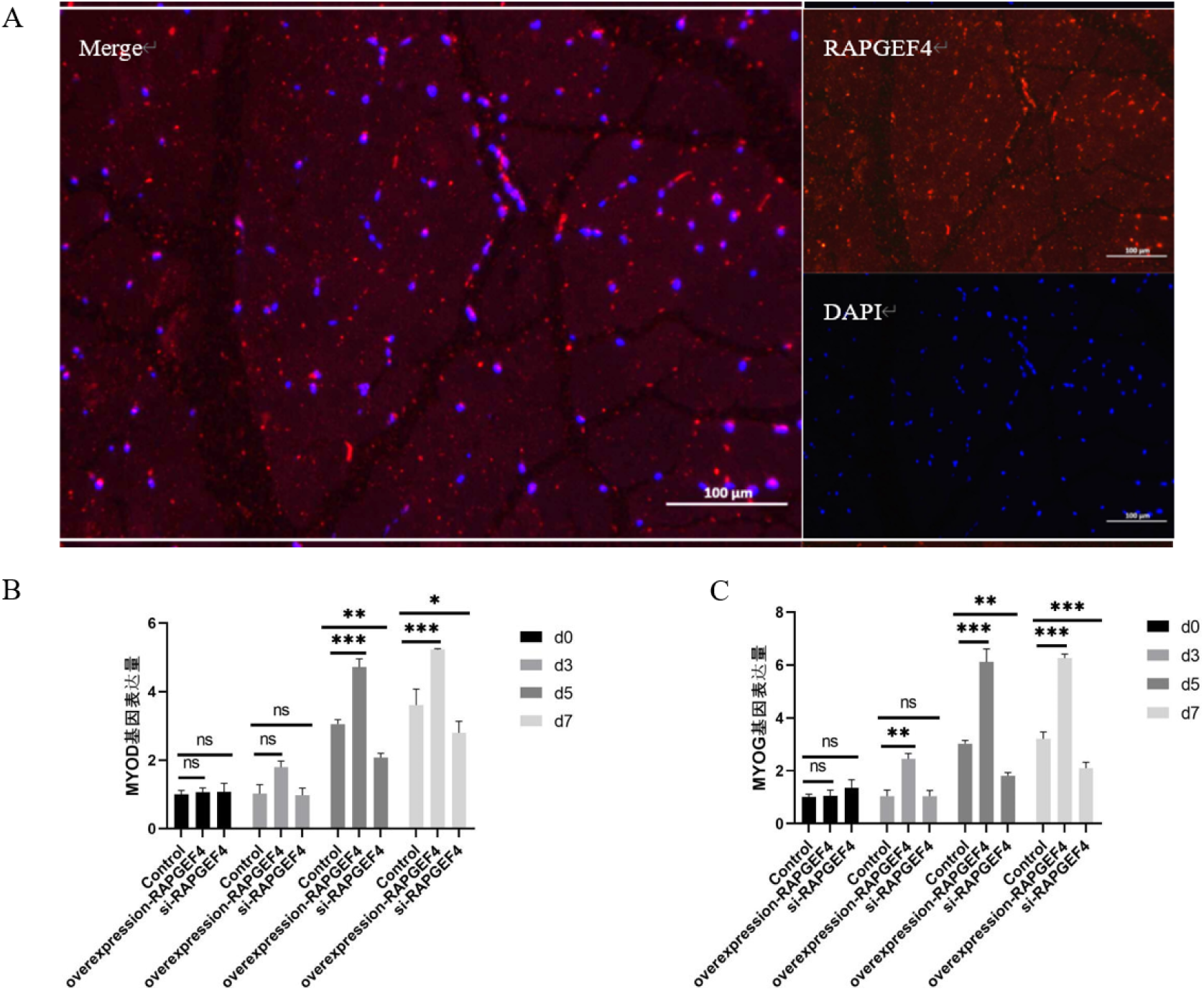
Effects of RAPGEF4 on C2C12 cell myogenic differentiation. a, RNA in situ hybridization verifying RAPGEF4 gene expression in the longissimus dorsi muscle tissue of Sunit Bactrian camels.saralab, Changes in expression levels of the myogenic marker gene MYOD at 0, 3, 5, and 7 days of myogenic induction differentiation in C2C12 cells (200× magnification) after RAPGEF4 gene overexpression and interference.saralac, Changes in expression levels of the myogenic marker gene MYOG at 0, 3, 5, and 7 days of myogenic induction differentiation in C2C12 cells (200× magnification) after RAPGEF4 gene overexpression and interference.

**Fig. 9.**
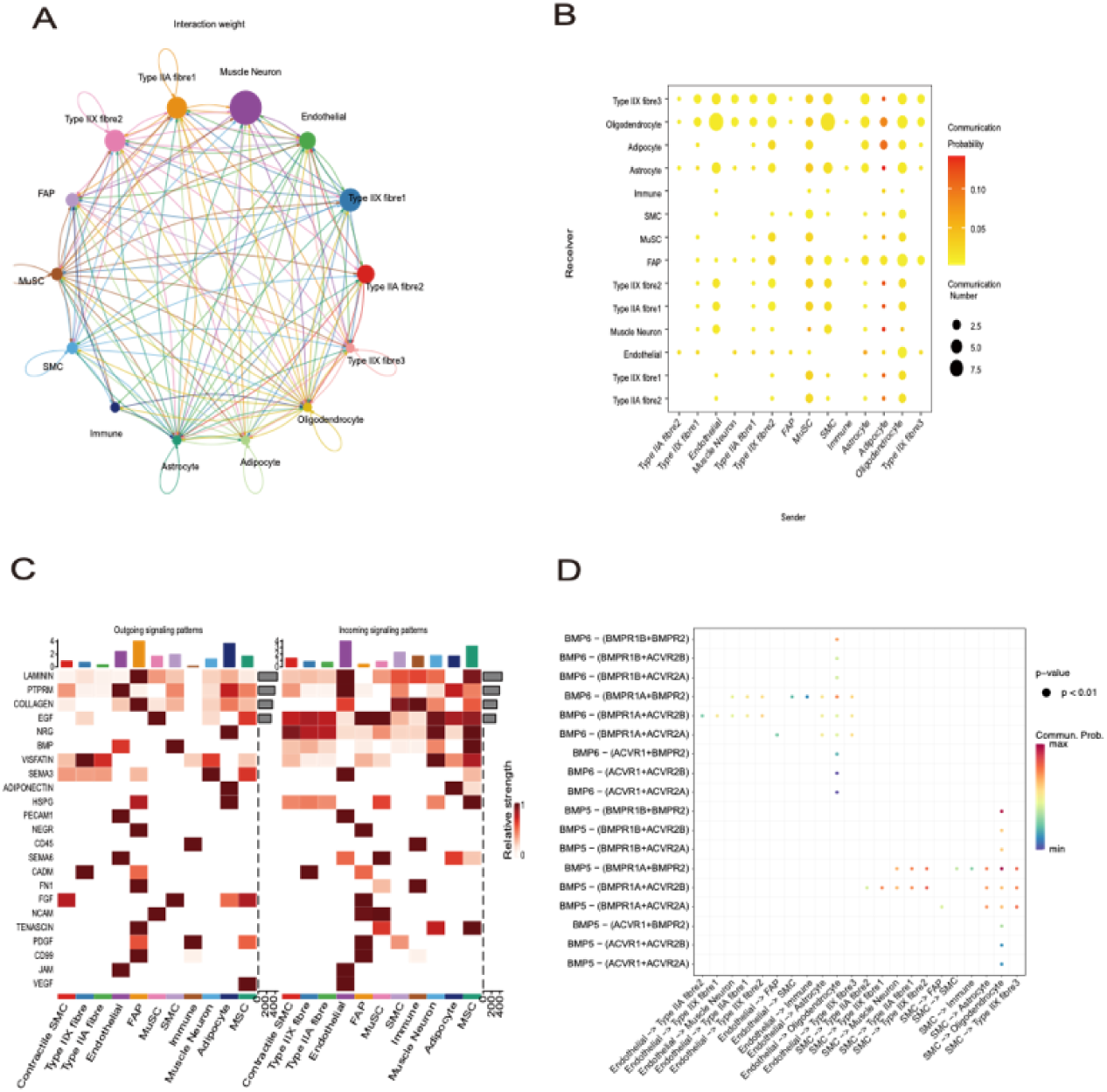
Analysis of intercellular communication patterns in longissimus dorsi muscle and exploration of the central role of the BMP pathway. a, Circular plot showing intercellular communication patterns in longissimus dorsi muscle. Line width represents the frequency of ligand-receptor interactions between two cell types, while circle size indicates the total number of interactions within each cell type. Different colours identify the signal-sending cell types.saralab, Dot plot detailing the number of significant interactions between each cell type pair. Circle size and colour correspond to the number of interactions and their probability, respectively.saralac, Depiction of major biological pathways in longissimus dorsi muscle and the intensity of signal output or input for each cell type, providing a comprehensive view of intercellular signalling strength and direction.saralad, Dot plot showing key ligand-receptor interaction pairs in the BMP pathway, with circle size and colour representing [incomplete sentence]

### 4.7 Impact of genetic selection signals on cell communication mechanisms

Intercellular communication is crucial for maintaining coordinated function, regulating development, homeostasis and tissue remodelling among individual cells within tissues. By leveraging cell type-specific transcriptional characteristics, this study delved into cell-cell interactions mediated by secreted and membrane-bound proteins, revealing synergies and regulatory mechanisms between different cell populations within tissues. This provides key insights into understanding the maintenance of dynamic equilibrium in organisms.

Given the widespread involvement of selection effects in gene regulation of Sunite camels, we hypothesized that selection effects might influence the expression activity of key ligand and receptor-related genes and participate in their interactions. To this end, we employed the CellChat algorithm to reveal the cell-cell communication landscape mediated by protein-ligand and receptor interactions in the longissimus dorsi muscle. Among 14 cell groups, we identified important ligand-receptor pairs and constructed a communication network between all cell types.

Notably, skeletal muscle stem cells exhibited frequent signalling with other cell types in this network, particularly through the BTC-ERBB4 ligand-receptor interaction in the EGF pathway, mediating signal transduction from skeletal muscle stem cells to various muscle fibre cells. This aligns with the finding of skeletal muscle stem cells as a differentiation starting point towards muscle fibre cells, highlighting their importance as a primary communication ‘hub’ in the muscle cell population.

Furthermore, SNP and indel sites altering element activity found in the C2C12 cell line may regulate the physiological modulation of skeletal muscle stem cells, thereby regulating the entire skeletal muscle population through cell communication.

In exploring other important cell pathways, we found that ACVR2A and ACVR1, two receptor genes in the BMP pathway, were affected by selection effects. Cell communication studies of the BMP pathway revealed their important roles in BMP6 pathway transmission from endothelial to oligodendrocyte cells and BMP5 pathway transmission from SMC to oligodendrocyte cells. This finding suggests that traits related to skeletal muscle and bone formation may have been regulated by selection effects during evolution.

These results not only highlight the crucial role of selection effects in cell communication but also provide new insights into understanding how these processes affect tissue function and organismal adaptability.

## 5 Discussion

This study, through the integration of multi-omics data including genome resequencing, epigenomics, and single-cell transcriptomics, comprehensively reveals the regulatory network and adaptive evolution mechanisms of the Bactrian camel genome for the first time. Our findings not only deepen the understanding of this important livestock species but also provide a solid scientific foundation for its genetic improvement and sustainable utilization.

By comparing the genomes of wild and domesticated Bactrian camels, we revealed significant genetic differences and genomic characteristics. These differences reflect the combined effects of natural and artificial selection, offering new perspectives on how Bactrian camels adapt to extreme environments and meet human needs. Significant genetic differentiation exists between wild and domesticated Bactrian camels, reflecting long-term evolutionary separation and different selection pressures. Mitochondrial genome analysis shows evidence of purifying selection between the two, implying conservation of certain gene functions during evolution^[55]^.

Genomic analysis revealed genetic variations in Bactrian camels adapting to specific environments. These variations likely originate from natural selection, enabling Bactrian camels to survive in extreme desert environments. The study identified a series of domestication-related genes that underwent positive selection. These genes may be associated with camels’ adaptation to human management, changes in behavioural characteristics, and adaptation to new diets and environments^[56]^.

Compared to previous studies^[57,58]^, our analysis is more comprehensive, including not only a larger sample size but also integrating multiple omics datasets, thus providing more detailed and reliable results.

By integrating epigenomic, transcriptomic, and single-cell data, we identified tissue-specific genes and regulatory elements in Bactrian camels, as well as genes under positive selection. These findings provide crucial clues for understanding the molecular mechanisms underlying complex traits in Bactrian camels. Notably, we constructed regulatory pathways from SNPs to regulatory elements to genes for the first time, filling a significant gap in Bactrian camel genomic research and guiding future functional validation experiments.

Our findings enrich the existing understanding of Bactrian camel muscle cell characteristics and their causal relationships with corresponding phenotypes. We revealed how to construct transcription factor-element-gene (TF-Element-Gene) regulatory networks by modulating skeletal muscle cell spectra, transcriptome expression, and ligand-receptor interactions across cell types to identify genes and their regulatory mechanisms associated with muscle traits.

Our study also mapped the single-cell transcriptional landscape of Bactrian camel skeletal muscle for the first time, revealing cellular heterogeneity and genetic expression patterns at different developmental stages and functional states. This discovery not only aids in understanding the molecular mechanisms of muscle development but also provides potential molecular targets for improving meat production traits in Bactrian camels.

During mammalian development, dynamic changes in chromatin states are closely related to cell differentiation processes, reflecting the remodeling of gene regulatory networks^[59]^. By tracking the myoblast differentiation process, this study revealed significant correlations between changes in open chromatin regions and gene expression patterns.

We found that MAP3K20 maintains high expression levels in late pseudotemporal cells and shows significant expression in all relevant single-cell clusters except intramuscular fat cells. This expression pattern suggests that MAP3K20 may play an important role in Bactrian camel skeletal muscle development.

Integrated single-cell RNA sequencing and single-cell assay for transposase-accessible chromatin (ATAC) sequencing analysis revealed that the RAPGEF4 gene exhibits cell type-specific gene regulation patterns in Bactrian camel skeletal muscle. Transcriptome, RT-PCR, and qPCR analyses indicated that overexpression of RAPGEF4 can promote myoblast differentiation.

The widespread application of whole-genome resequencing and epigenomic technologies has facilitated the identification of highly differentiated genomic regions, becoming a crucial starting point for exploring the genetic basis of domestication. However, functional coding variants account for only a minority, making it challenging to fully explain observed selection signals^[60,61,62]^. These complex factors significantly increase the technical challenges of precisely locating potential causal variants and related genes.

Our study systematically analyzed gene expression and chromatin accessibility across multiple Bactrian camel tissues. Utilizing these rich functional annotation data, we developed a novel method that significantly enhanced our ability to decipher the genetic basis of Bactrian camel domestication and breeding. To our knowledge, this is the first study applying multi-tissue functional genomic annotations to investigate the impact of non-coding variants on phenotypic diversity in Bactrian camels.

The analytical framework established in this study is highly flexible and scalable, integrating chromatin accessibility and gene expression data, and can easily incorporate other types of functional genomic annotations. Notably, by introducing genome-wide chromosome three-dimensional structure data (such as Hi-C), we can more precisely interpret the relationships between distal regulatory elements and their target genes. This approach will significantly improve our accuracy and efficiency in identifying and assigning functional distal features, such as enhancers.

For example, using chromatin long-range interactions revealed by Hi-C data, we can more reliably associate specific regulatory elements with their regulated genes, thereby comprehensively understanding the complexity of gene regulatory networks. This integration strategy is not only applicable to Bactrian camels in this study but can also be extended to functional genomic research in other species, providing a powerful tool for elucidating the genetic basis of complex phenotypes.

This study reveals genetic differences and genomic features between wild and domesticated Bactrian camels through whole-genome resequencing of 35 individuals from the Inner Mongolian Sunite local population, combined with public data from 112 Asian Bactrian camels. We integrated multi-omics data, including ATAC-seq, RNA-seq, scRNA-seq and scATAC-seq, to identify tissue-specific genes, regulatory elements and genes under selection. These findings provide new perspectives on the adaptive evolution of Bactrian camels.

Notably, we found that selection effects on skeletal muscle traits involve not only mutations in coding regions but also SNPs and indels in regulatory elements. These sites may influence the survival ability and economic traits of Bactrian camels by altering the activity of cis-regulatory elements or their binding affinity with transcription factors.

Through single-cell transcriptome analysis, we revealed cellular heterogeneity and gene expression patterns during skeletal muscle development, providing new insights into the adaptive evolution and formation of economic traits in Bactrian camels.

This study enriches our understanding of the genomic structure and function of Bactrian camels, highlighting the important role of epigenetic regulation in animal adaptive evolution and the formation of economic traits. These findings provide a scientific basis for breeding and genetic resource conservation of Bactrian camels and other livestock, while offering important data support for future genetic improvement of Bactrian camels.

## 6 Author Contributions

## 8 Code Availability

## 9 Supplementary Information

